# A computational approach to identify possible symbiotic mechanisms between *Klebsiella* and the Mediterranean fly (*Ceratitis capitata*)

**DOI:** 10.1101/2021.02.13.431071

**Authors:** L. Augusto Franco, César Alberto Bravo Pariente

**Affiliations:** Programa de Pós-Graduação em Modelagem Computacional em Ciência e Tecnologia, Universidade Estadual de Santa Cruz, Ilhéus, Bahia, Brazil

**Keywords:** SIT, Medfly, *Klebsiella*, Reconstruction, Metabolic, Pathways

## Abstract

Nowadays, scientists develop multiple methods to control pests, like chemical control as the insecticides and biological control as the Sterile Insect Technique (SIT) to release infertile males produced by irradiation. This technique is complemented with research in the medfly’s *(Ceratitis capitata)* microbiota, this approach is based on the identification of molecular mechanisms from three *Klebsiella* strains using the reconstruction of metabolic pathways technique. We focus on the reconstruction of metabolic pathways involved in nitrogen metabolization because the bacteria are in charge of metabolizing the nitrogen compounds and make available for the medfly. We propose a pipeline to process the genome sequences the bacteria, reconstruct metabolic pathways and identify possible symbiont molecular mechanisms. The result of processing the biological data were ten pathways, six pathways produce L-Glutamate as final product, three pathways with ammonia product and one pathway was discarded because it was an internal pathway and don’t generate a metabolic product. Those pathways corresponding to L-Glutamate and ammonia metabolic products are part of nitrogen metabolism on *Klebsiella*, which shows that this metabolic process is one of the molecular mechanisms involved in symbiosis process between *Klebsiella* and the medfly. Finally, by the results of the sequences analysis, in the medfly’s metamorphose the L-Glutamate are used in the synthesis of new proteins and the production of energy. The final consideration suggest that the pipeline propose can be used as first step to identify molecular mechanisms that can improve the production of industrial the medfly.

## Introduction

The Sterile Insect Technique (SIT) is one of the most insect pest control techniques used. This technique is an environmentally friendly method that consists of mass-rearing and sterilization by irradiation of the male insects to control the size of the insect population. The SIT technique is used in the Medfly *(Ceratitis capitata)* to release sterile males over defined areas, where the male found wild females to copulate, resulting in no offspring reducing the pest population (Klassen et al., 2005). As well as, in the process of improving the biological control of this organism, new approaches are applying. For example, research of the midgut microbiota from *C. capitata* and identifying potentially symbiotic bacteria that have a positive effect on the development and fitness of the sterile medfly.

It is known that bacteria of the *Klebsiella* genus plays a crucial role on the nitrogen metabolization that is used by the medflies in the digestive process (Ami et al., 2009). Also, the symbiotic relationship between *Klebsiella* and the medfly is present at the metamorphosis process of the medfly. This biochemical process involves the protein digestion between stages of metamorphosis which produce free amino acids that are used in the synthesis of new proteins in the following metamorphosis processes and also can be oxidated to enter in the tricarboxylic acid cycle (TCA) (Nestel et al., 2003) to produce immediate energy for the metamorphosis process. Consequently, these free amino acids are nitrogen compounds, that *Klebsiella* can metabolize and make available for the medfly.

There are several molecular mechanisms in *Klebsiella* spp. that can improve the behavior of the irradiated medfly males. We propose a pipeline to reconstruct metabolic pathways of three *Klebsiella* strains, compare them and identify molecular mechanism that can be object of metabolic or genetic engineering. The principal ideas are also present the *Klebsiella* bacteria as a possible medfly symbiont to improve the biological control of the medfly. In the context of computational biology, be can use genomes information to reconstruct metabolic pathways. The metabolic pathways describes systematic functions that occur through metabolites (cellular components involved in metabolism), and these components are connected to generate metabolic product (Faust et al., 2011). The metabolization of nitrogen in *Klebsiella* spp. was the target for reconstruct the metabolic pathway involved in this metabolic system. These pathways were analyzed by the stoichiometric modeling of the reaction and the Elementary Modes (EM) analysis approach (Feist, et al., 2008).The EM analyze all the possible flows (pathways) involved in the proposed metabolic system of nitrogen metabolization (Schilling et al, 2000).

In order to fulfill the objective of this paper, three genomes of *Klebsiella* bacteria were processed with computational tools to identify molecular mechanism on *Klebsiella* that can improve the SIT technique on *C. capitata.* This study was complemented using stoichiometric modeling of reactions involved in nitrogen metabolization and analyzing them with EM approach. Those results were compared with metamorphosis data of medfly.

## Materials and methods

In order to reconstruct the metabolic pathways of three *Klebsiella* bacteria genomes we propose a computational pipeline (Fig. 1) to manipulate biological data of sequencing raw results and a description of how the pipeline can be used to reconstruct metabolic pathways from genome-scale. In this case, we use this technique to identify molecular mechanisms present in *Klebsiella* as a potential tool to improve SIT technique in *C. capitata.*

**Fig. 1.**
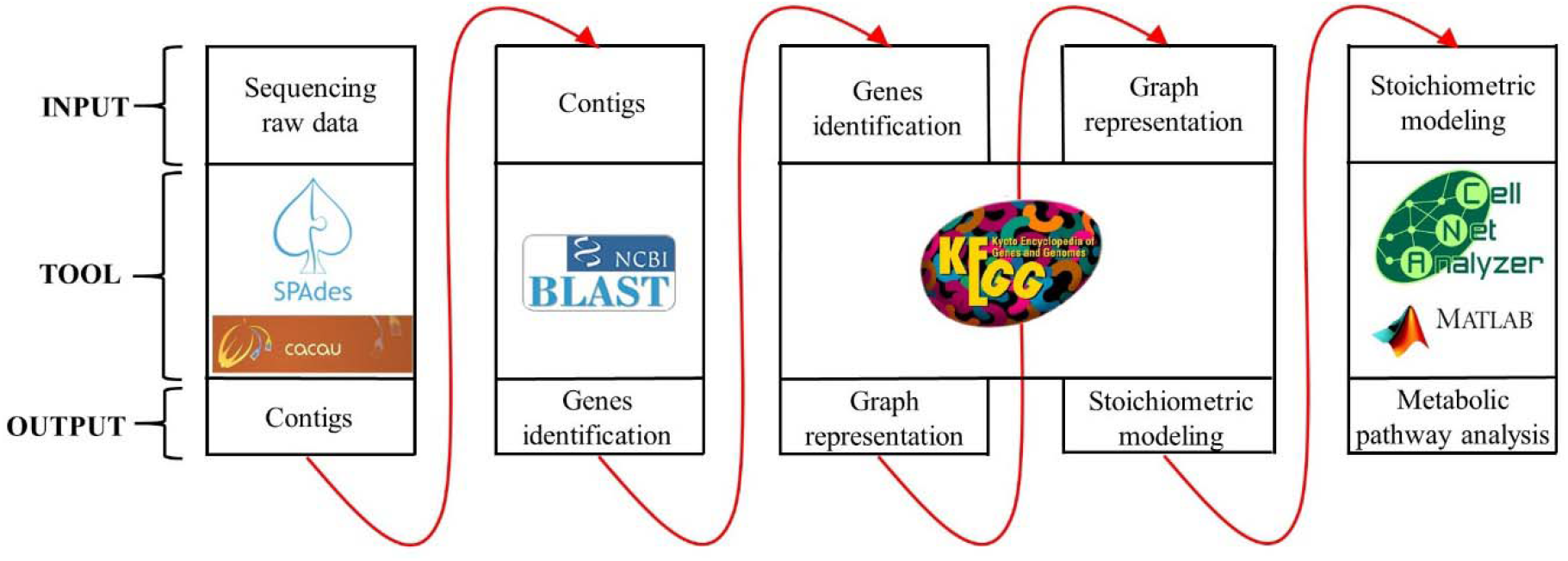
Pipeline proposed to metabolic pathway reconstruction from genome-scale. This figure shows the data needed for input, computational tools and the output of each step in this process.

### Data availability

The biological data was provided from a previous research conducted at Universidad del Valle de Guatemala, Guatemala (Figueroa & Navarrete, 2018). The products of this research were three genome sequencing results of *Klebsiella* strains. The Whole-Genome Sequencing (WGS) projects is deposited on GenBank under the accession numbers (JACOEB000000000, JACOEC00000000 and JACOED000000000). Raw reads are deposited in the SRA under the accession numbers (SRR12450864, SRR12450865 and SRR12450866). These samples are part of BioProject accession number (PRJNA656584). Additionally, we work with genes sequences obtained from the National Center for Biotechnology Information (NCBI), the European Bioinformatics Institute (EBI) and the Kyoto Encyclopedia of Genes and Genomes (KEGG).

### Contigs assembly

The contigs assembly was performed in the first step of the pipeline with the supercomputer “Centro de Armazenamento de dados e Computação Avançada da UESC” (CACAU, by its acronym in Portuguese) with this description: Bull novascale, 20 nodes R422-E1, two processors Intel^®^Xeon^®^ E5430 @ 2.66GHz QuadCore, 16 GB memory, one InfiniBand Board, one Gigabit Ethernet Board (Total: 160 Cores and 320 GB memory). The assembly was performed by the SPAdes genome assembler (v 3.13.0) (Bankevich et al., 2012) with paired reads (2 x 150 bp) results of the three *Klebsiella* strains at CACAU.

### Identification of symbiotic genes

In the second step of the pipeline the contigs obtained were aligned against sequences of symbiotic genes involved in nitrogen metabolization (Supplementary Data Table S1 and S2), obtained from NCBI, EMBL-EBI and KEGG databases. These alignments were performed by BLAST tool (v2.8.1) (Altschul, et al., 1990). This process was performed for the contigs of the three *Klebsiella* strains.

### Identification of metabolic pathways

The third step of the process was performed processing the genes identified for the three strains of *Klebsiella* in KEGG database (v2019) (Ogata et al., 1999). The entry data of this database were the genes identified by the BLAST alignment. The result was a graph representing the metabolic pathways where the genes products were involved. However, a subgraph was extracted containing only the information related to genes products involved in the nitrogen metabolization of *Klebsiella* bacteria.

### Stoichiometric modeling of metabolic pathways

Considering the subgraph extracted for each strain of bacteria, in the fourth step of the process, we performed the stoichiometric modeling of these metabolic pathways. In this stage the metabolic pathways were mathematically represented by a stoichiometric matrix (*S*) (Maarleveld et al., 2013). This matrix was built with the data of the number of metabolites (nodes) in the subgraph and the stoichiometric coefficients for each reaction in the metabolic pathways extracted for KEGG database. The matrix *S* = *m* x *n* was built according to number of metabolites (*m*) and reactions (*n*) involved in nitrogen metabolization in *Klebsiella* strains.

### Analysis of metabolic pathways using CellNetAnalyzer (CNA)

In fifth step of the pipeline the analysis of metabolic pathways was performed using CellNetAnalyzer (v2019.1) (Klamt et al., 2007) in MATLAB (v8.4) interface. This tool allows us to adapt the metabolic pathway analysis in our data performing the Elementary Modes analysis. This analysis calculates all the possible solutions for pathways of incoming exchange metabolites to output metabolites in the nitrogen metabolization subgraph (Trinh et al., 2009). For our model, we based our research on the symbiotic genes identified on the BLAST, 11 metabolites were considered; in addition, twelve reactions (internal paths), two input metabolites (Nitrate and S-Allantoin) and two output metabolites (Ammonia and L-Glutamate).

## Results

### Contigs Assembly

The SPAdes tool produces assemblies contigs for each strain of *Klebsiella* bacteria as show in Table 1. All the contigs generated were indispensable due the sequences of DNA contained in each one.

**Table 1.**
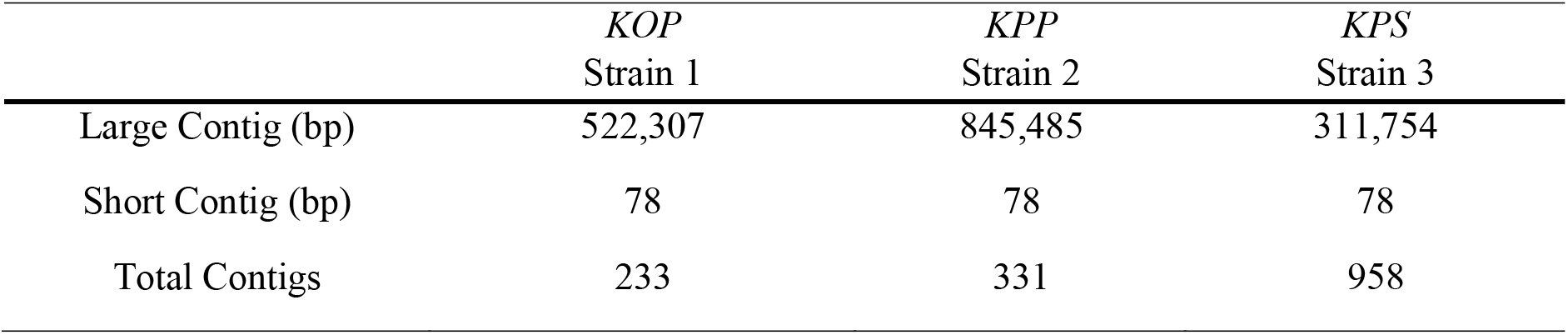
Summary of contigs obtained for *Klebsiella* bacteria.

### Identification of potentially symbiotic genes

The genes identified (Supplementary Data Tables S1 and S2) were present in the contigs obtained for each strain of bacteria. These genes are involved in nitrogen metabolization in *Klebsiella.* Additionally, is known that they have a potentially symbiotic function in *C. capitata* (Aharon et al., 2013). These genes are classified in groups according to the functions: genes for nitrogen fixation, nitrogen assimilation and nitrogen regulation. The other group of genes are involved in the synthesis of enzymes involved in reactions of the nitrogen metabolization in *Klebsiella.*

### Identification of metabolic pathways

These genes are involved in the synthesis of enzymes or metabolites present in nitrogen metabolism of Klebsiella bacteria. According to graph theory, we represent 11 metabolites (Fig. 2A) and coded from 0 to 10 according to the node’s numeration. Considering each metabolite, our results show the subgraph obtained (Fig. 2B), it was designed to represent the metabolic pathways of nitrogen metabolization in Klebsiella bacteria.

### Stoichiometric modeling

In this stage we construct the stoichiometric matrix (*S*) (see Supplementary Data Fig. S1 for details), constructed with 16 columns corresponding to the number of reactions involved in the subgraph previously obtained. In addition, 11 lines corresponding to the metabolites involved in nitrogen metabolization reactions. The values present in this matrix correspond to stoichiometric values for each reaction, “0” represents that the metabolite is not used in the reaction, a negative value is a metabolite consumed and a positive value is a metabolite produced.

### Analysis of metabolic pathways

The metabolic pathways identified and the data provided by stoichiometric modeling were used to calculate the Elementary Modes analysis in our metabolic pathway reconstructions. The results were all the combinations of reactions that form one metabolic pathway (the graphical representations of these EM are provided in the Supplementary Data Fig. S2 and Fig. S3). We obtained ten Elementary Modes represented in *P* matrix (Fig. 3), each EM was codified as *p_i_* where *i* represent the number of EM. We represent metabolites as nodes and reactions as edges. The results present, that we obtained six EM that the final product was L-Glutamate, three EM with Ammonia and one EM that was no useful for our research.

**Fig. 3.**
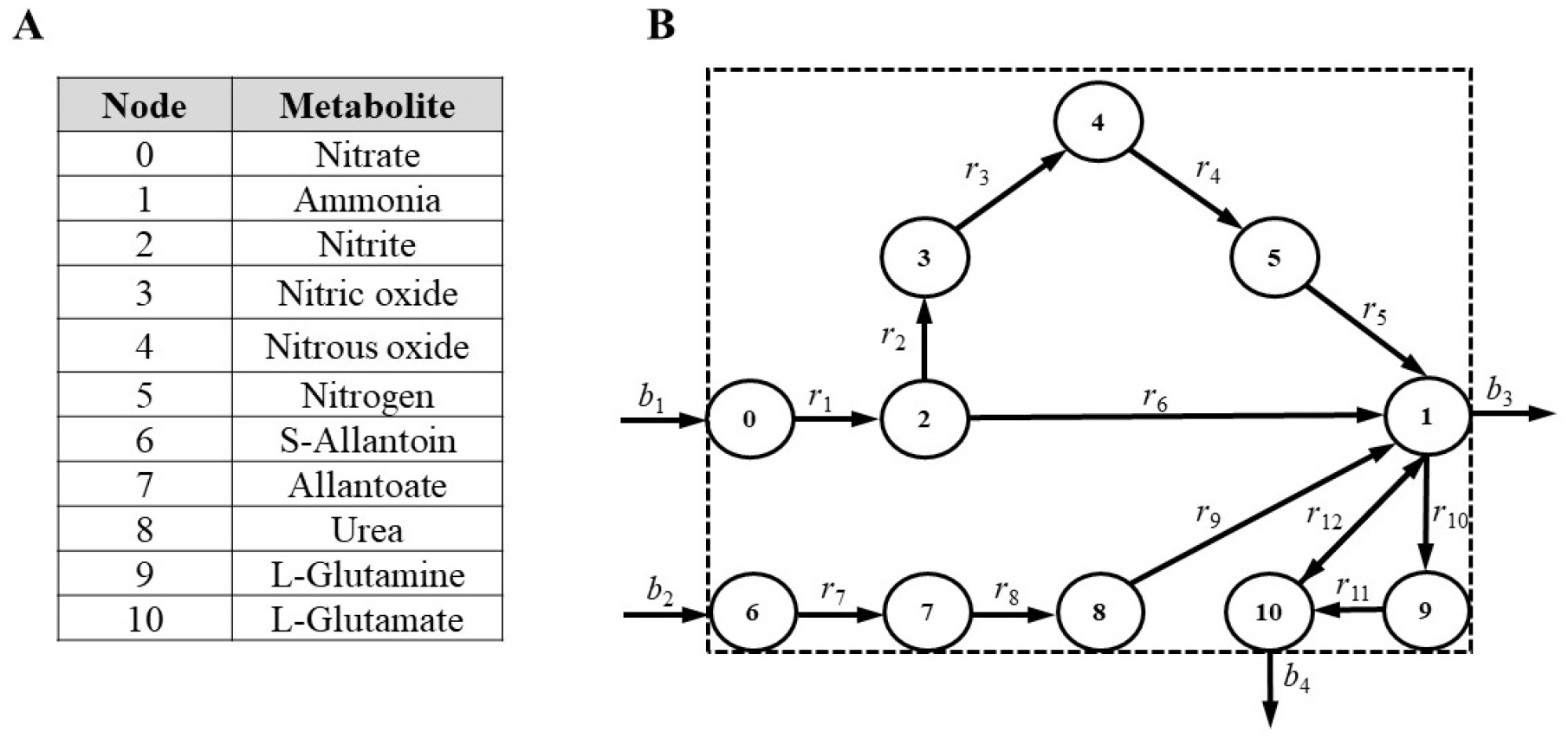
Identification of nitrogen metabolization pathways in *Klebsiella* strains based on metabolites and reactions involved. We represent metabolites as nodes and reactions as edges, *bi* exchange reactions and *ri* internal reactions. **A**) Nodes codifying of metabolites. **B**) Subgraph of nitrogen metabolization in *Klebsiella* bacteria.

**Fig. 3.**
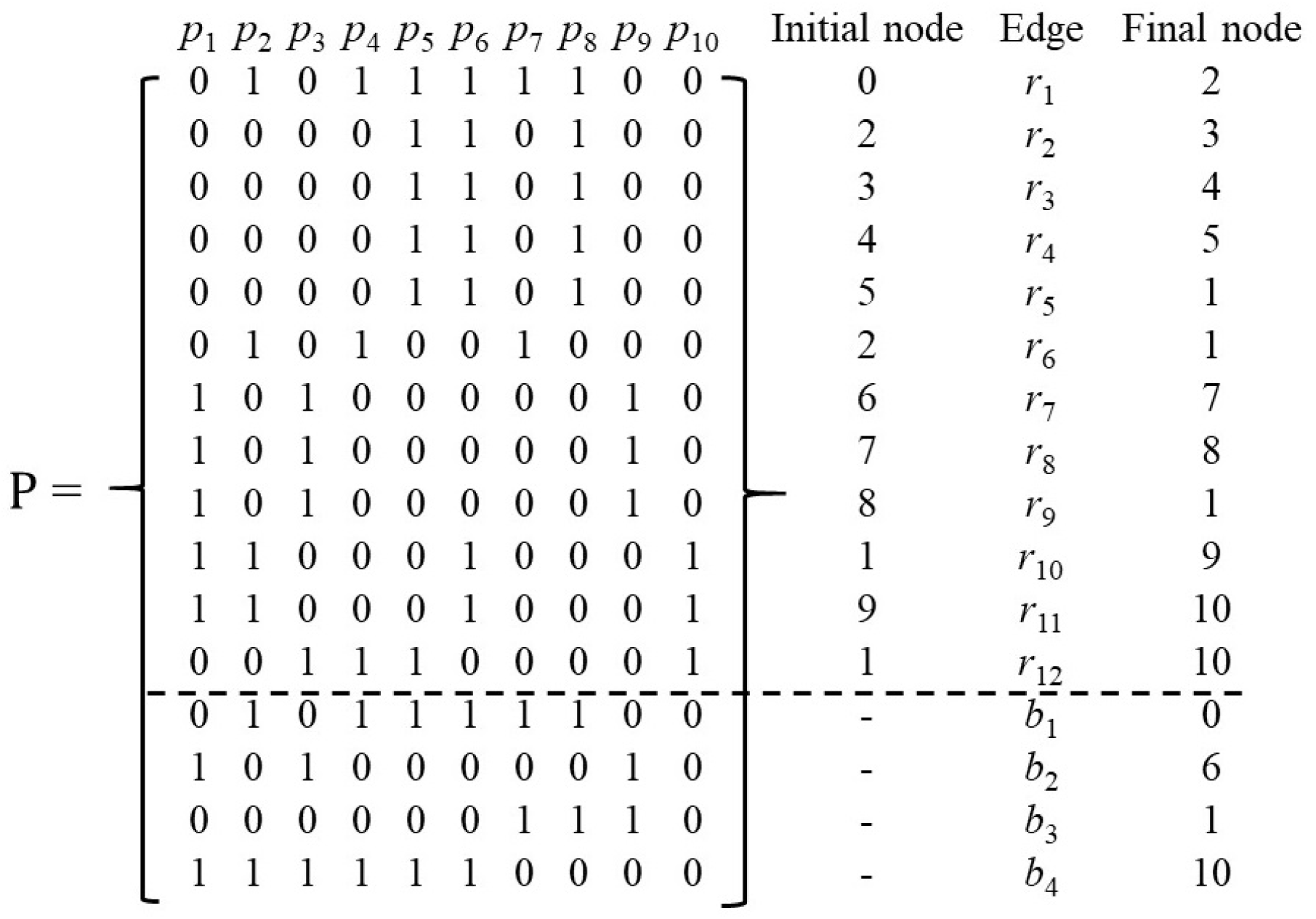
Matrix representation of EM obtained from metabolic pathway analysis involved in nitrogen metabolization in *Klebsiella* bacteria. This matrix shows the summary of metabolic pathway analysis by Elementary Modes. Each column of the matrix represent one EM obtained (*p*_i_). Also, we show the initial and final node and the reactions involved.

## Discussion

The biological control of *Ceratitis capitata* using SIT technique produces adult males with unviable spermatozoa in order to reduce the population of medfly in the environment. Therefore, the SIT technique is constantly improved, and one approach used is the addition of bacteria strains in medflies diet. These bacteria are known to be potentially symbiotic with the Medfly. One of the most present bacteria in Medfly microbiota is *Klebsiella*, which can metabolize nitrogenous compounds that medfly needs for its development (Aharon et al., 2013). The metabolization of nitrogen in these bacteria can be identified with the application of the technique of reconstruction of metabolic pathway from the genome of *Klebsiella* bacteria identified at the medfly’s production facility.

The three sequencing data obtained from Figueroa & Navarrete, 2018 (Unpublised data), were our starting point. These three sequencing results were processed with the pipeline proposed to reconstruct metabolic pathways from genome scale. These assembly results show difference in the amount of contigs generated for each strain, and those differences were observed due we work with biological data of different strains of *Klebsiella* bacteria. Even though the difference may be related to the presence/absence of certain genetic sequences or the origin of each isolate.

On the other hand, genes involved in nitrogen metabolization in the three *Klebsiella* bacteria were identified with BLAST tool (Altschul, et al., 1990). These genes have a symbiotic relationship with *C. capitata,* because they are involved in fixation, assimilation and regulation of nitrogen in *Klebsiella.* In the case of these genes, we identified a difference between the genes involved in nitrogen fixation for the three strains of bacteria. The difference is due because among species most genes are conserved like *Klebsiella*. Also, the genes involved in nitrogen metabolization are crucial because most of the diet of medfly is poor in nitrogen compounds (Slansky Jr., 1985; Waldbauer & Friendman, 1991) and a medfly needs the symbiotic relationship with *Klebsiella* bacteria to obtain those nitrogen compounds, like L-Glutamate and glutamine. These compounds can be used in the synthesis of new proteins in metamorphosis process of *C. capitata.*

The results of metabolic pathway analysis of the three strains of *Klebsiella* were similar, because they show the same reactions and metabolites involved for each strain and the same EM generated. Consequently, we proposed a construction of a criterion based on the energy consumption that occurs in medfly metamorphosis. The first approach to construct a criterion was based on the fixed output metabolites in Elementary modes analysis. Ammonia is a toxic nitrogenous compound for organisms, so to reuse it in the cycle allows to synthesize other important molecules for bacteria metabolism. One of these molecules is L-Glutamate, it is an amino acid of interest in symbioses with medfly, because is used to perform the synthesis of new proteins and ATP at the stages of the metamorphosis of medfly.

In metamorphosis stages the medfly needs energy in multiple forms to accomplish the transformation because the medfly suffers biochemical process necessary for development. So, we focused in proteins as an energy source in medfly metamorphosis. According to (Nestel et al., 2003) there exists digestion of proteins in the process of metamorphosis, with the production of free amino acids and energy flux in the organism. Therefore, the medfly metamorphosis occurs in four stages, from egg, to larval, to pupal and finally to adult. In the larval stage the larval cuticle protein is present and the amino acids that form this protein are free in the medium. Additionally, the pupal cuticle protein begins to synthesis with the free amino acid’s product of larval cuticle protein digestion. Our results shows that L-glutamate is one of the major products of *Klebsiella* nitrogen metabolization. The metabolization of this amino acid can be use to synthesize larval and pupal cuticle, but most important, this amino acid can be oxidated and be used at the TCA to produce energy.

## Conclusions

We identified possible metabolic pathways involved in the process of symbiosis between *Klebsiella* bacteria and *Ceratitis capitata* using our pipeline. Those metabolic pathways were compared between the three *Klebsiella* strains. We propose the nitrogen metabolization mechanisms as a possible target of metabolic and genetic engineering in *Klebsiella* that can be added in diet of *C. capitata* to improve the production by SIT technique. The *Klebsiella* bacteria can metabolize free amino acids to use them for protein synthesis and energy production, especially the L-Glutamate that can act as a building block for proteins and can be oxidated to α-ketoglutarate, enter to the TCA and produce ATP. This ATP molecules will be crucial at the metamorphosis process of the medfly. If the sterile males of *C. capitata* can survive the metamorphosis process they will be near to complete the mission of: reduce the medfly population.

## Declarations

Not applicable

## Ethics approval

Not applicable

## Consent to participate

Not applicable

## Consent for publication

Not applicable

## Availability of data and material

### Code availability

Not applicable

### Authors Contributions

Not applicable

## Acknowledgements

We thank to the Universidad del Valle de Guatemala and Programa MOSCAMED – Guatemala for the data proportionated. We thank the Coordenação de Aperfeiçoamento de Pessoal de Nível Superior (CAPES) and the Programa de Pós-Graduação em Modelagem Computacional em Ciência e Tecnologia (Universidade Estadual de Santa Cruz-UESC) for the master’s scholarship. Thanks to the staff of Nucleo de Biologia Computacional e Gestão de Informações Biotecnológicas (NBCGIB) for the space and help provided at the time when the project was developed.

## References

Aharon, Y., Pasternak, Z., Yosef, M. Ben, Behar, A., Lauzon, C., Yuval, B., & Jurkevitch, E. (2013). Phylogenetic, metabolic, and taxonomic diversities shape mediterranean fruit fly microbiotas during ontogeny. Applied and Environmental Microbiology, 79(1), 303–313. https://doi.org/10.1128/AEM.02761-12

Altschul, S. F., Gish, W., Miller, W., Myers, E. W., & Lipman, D. J. (1990). Basic local alignment search tool. Journal of Molecular Biology, 215(3), 403–410. https://doi.org/10.1016/S0022-2836(05)80360-2

Ami, E. Ben, Yuval, B., & Jurkevitch, E. (2009). Manipulation of the microbiota of mass-reared Mediterranean fruit flies Ceratitis capitata (Diptera: Tephritidae) improves sterile male sexual performance. TheISME Journal, 4(1), 28–37. https://doi.org/10.1038/ismej.2009.82

Bankevich, A., Kulikov, A. S., Prjibelski, A. D., Tesler, G., Vyahhi, N., Sirotkin, A. V., Pham, S., Dvorkin, M., Pevzner, P. A., Bankevich, A., Nikolenko, S. I., Pyshkin, A. V., Nurk, S., Gurevich, A. A., Antipov, D., Alekseyev, M. A., & Lesin, V. M. (2012). SPAdes: A New Genome Assembly Algorithm and Its Applications to Single-Cell Sequencing. Journal of Computational Biology, 19(5), 455–477. https://doi.org/10.1089/cmb.2012.0021

Faust, K., Croes, D., & Helden, J. Van. (2011). Prediction of metabolic pathways from genome-scale metabolic networks. BioSystems, 105(2), 109–121. https://doi.org/10.1016/j.biosystems.2011.05.004

Feist, A. M., Herrgård, M. J., Thiele, I., Reed, J. L., & Palsson, B. Ø. (2008). Reconstruction of biochemical networks in microorganisms. Nature Reviews. Microbiology, 7, 129–143. https://doi.org/10.1038/nrmicro1949

Figueroa, N., & Navarrete, C. (2018). Caracterización genética de cepas de Klebsiella spp. aislada de ambientes asociados a Ceratitis capitata (Weidemann), por medio de secuenciación de su genoma, extracción de plásmidos y Rep-PCR. Universidad del Valle de Guatemala.

Klamt, S., Saez-Rodriguez, J., & Gilles, E. D. (2007). Structural and functional analyzis of cellular network with CellNetAnalyzer. BMC Systems Biology, 2(1), 13. https://doi.org/10.1186/1752-0509-2-104

Klassen, W., Curtis, C. F., Dyck, V. A., Hendrichs, J., & Robinson, A. S. (2005). Sterile Insect Technique Principles and Practice in Area-Wide Integrated Pest Management. Springer.

Maarleveld, T. R., Khandelwal, R. A., Olivier, B. G., Teusink, B., & Bruggeman, F. J. (2013). Basic concepts and principles of stoichiometric modeling of metabolic networks. Biotechnology Journal, 8(9), 997–1008. https://doi.org/10.1002/biot.201200291

Nestel, A. D., Tolmasky, D., Rabossi, A., Luis, A., Nestel, D., Tolmasky, D., & Rabossi, A. (2003). Lipid, Carbohydrates and Protein Patterns During Metamorphosis of the Mediterranean Fruit Fly, Ceratitis capitata (Diptera: Tephritidae) Published By ?: Entomological Society of America Lipid, Carbohydrates and Protein Patterns During Metamorphosis o. Entomological Society of America, 96(3), 237–244.

Ogata, H., Goto, S., Sato, K., Fujibuchi, W., Bono, H., & Kanehisa, M. (1999). KEGG: Kyoto encyclopedia of genes and genomes. Nucleic Acids Research, 27(1), 29–34. https://doi.org/10.1093/nar/27.1.29

Schilling, C. H., Letscher, D., & Palsson, B. O. (2000). Theory for the systemic definition of metabolic pathways and their use in interpreting metabolic function from a pathway-oriented perspective. Journal of Theoretical Biology, 203(3), 229–248. https://doi.org/10.1006/jtbi.2000.1073

Slansky Jr., F. (1985). Food utilization by insects: interpretation of observed differences between dry weight and energy efficiencies. Entomologia Experimentalis et Applicata, 39(1), 47–60.

Trinh, C. T., Wlaschin, A., & Srienc, F. (2009). Elementary Mode Analysis: A Useful Metabolic Pathway Analysis Tool for Characterizing Cellular Metabolism. Applied Microbiology, 81(5), 813–826. https://doi.org/10.1007/s00253-008-1770-1.Elementary

Waldbauer, G. P., & Friendman, Sa. R. of E. (1991). Self-Selection of Optimal Diets by Insects. Annual Review of Entomology, 36(1), 43–63.

