## Supplemental Files for "A computational approach to identify possible symbiotic mechanisms between *Klebsiella* and the Mediterranean fly (*Ceratitis capitata*)"

Supplementary data:

Summary information of genes with potentially symbiotic effect between *Klebsiella* spp., and *Ceratitis capitata*. In Table S1 we described the identified genes in KOP strain. In Table S2 we described the identified genes for KPP *and* KPS.

| **Gene** | **Function** | **Length (bp)** | **Accession number** | **Organism** |
| --- | --- | --- | --- | --- |
| *nifH* | Nitrogen fixation | 263 | KJ940124.1 | *Klebsiella* sp. |
| *nifD* |  | 1523 | Y00316.1 | *Klebsiella* sp. |
| *nifK* |  | 1557 | CAA29588.1 | *Klebsiella* sp. |
| *allS* | Nitrogen assimilation | 738 | MF417539.1 | *Klebsiella* sp. |
| *allC* |  | 2516 | SAMEA2273639_02901 | *K. Oxytoca* |
| *ureG* |  | 618 | 29381131 | *Klebsiella oxytoca* |
| *glnA* | Glutamine sintetase synthesis | 928 | LC011557.1 | *Klebsiella* sp. |
| *gltB* | Glutamate synthesis | 7930 | AY035435.1 | *Klebsiella* sp. |
| *gluD* | Glutamate dehydrogenase synthesis | 1382 | 29379502 | *Klebsiella oxytoca* |
| *ntrA* | Nitrogen regulation | 1935 | X03147.1 | *Klebsiella* sp. |
| *ntrB* |  | 1408 | X03146.1 | *Klebsiella* sp. |
| *ntrC* |  | 1575 | X02617.1 | *Klebsiella* sp. |
| *narI* | Nitrate reductase | 681 | KONIH1_14050 | *Klebsiella oxytoca* |
| *narH* |  | 1545 | KONIH1_14060 | *Klebsiella oxytoca* |
| *narZ* |  | 3741 | KONIH1_14070 | *Klebsiella oxytoca* |
| *norBC* | Nitric oxide reductase | 2734 | KONIH1_22945 | *Klebsiella oxytoca* |
| *nirK* | Nitrite reductase | 2874 | AAA25099.1 | *Klebsiella oxytoca* |

**Supplementary Table S1.** Identified genes from contigs of KOP strain.

**Supplementary Table S2.** Identified genes from contigs of KPP and KPS

| **Gene** | **Function** | **Length (bp)** | **Accession number** | **Organism** |
| --- | --- | --- | --- | --- |
| *nifU* | Nitrogen fixation | 387 | KPN_02861 | *Klebsiella pneumoniae* |
| *hpxB* | Nitrogen assimilation | 933 | KPN_01787 | *Klebsiella pneumoniae* |
| *allC* |  | 1260 | KPN_01761 | *Klebsiella pneumoniae* |
| *ureG* |  | 618 | 29381131 | *Klebsiella oxytoca* |
| *glnA* | Glutamine sintetase synthesis | 928 | LC011557.1 | *Klebsiella* sp. |
| *gltB* | Glutamate synthesis | 7930 | AY035435.1 | *Klebsiella* sp. |
| *gluD* | Glutamate dehydrogenase synthesis | 1382 | 29379502 | *Klebsiella oxytoca* |
| *ntrA* | Nitrogen regulation | 1935 | X03147.1 | *Klebsiella* sp. |
| *ntrB* |  | 1408 | X03146.1 | *Klebsiella* sp. |
| *ntrC* |  | 1575 | X02617.1 | *Klebsiella* sp. |
| *narI* | Nitrate reductase | 681 | KONIH1_14050 | *Klebsiella oxytoca* |
| *narH* |  | 1545 | KONIH1_14060 | *Klebsiella oxytoca* |
| *narZ* |  | 3741 | KONIH1_14070 | *Klebsiella oxytoca* |
| *norBC* | Nitric oxide reductase | 2734 | KONIH1_22945 | *Klebsiella oxytoca* |
| *nirK* | Nitrite reductase | 2874 | AAA25099.1 | *Klebsiella oxytoca* |

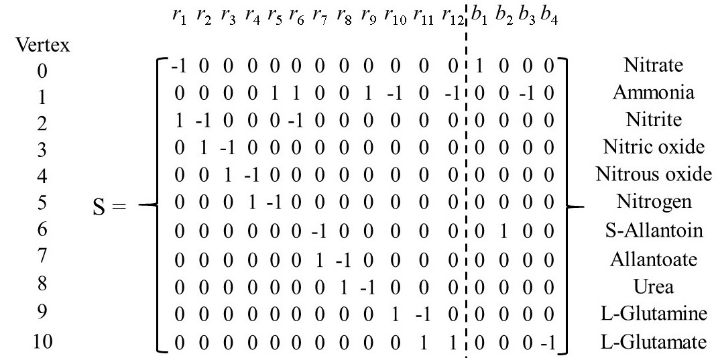

**Supplementary Fig. S1.** Stoichiometric matrix of metabolic pathways involved in nitrogen metabolization in *Klebsiella* bacteria. This matrix shows the metabolites and reactions that occur in nitrogen metabolization in *Klebsiella* bacteria.

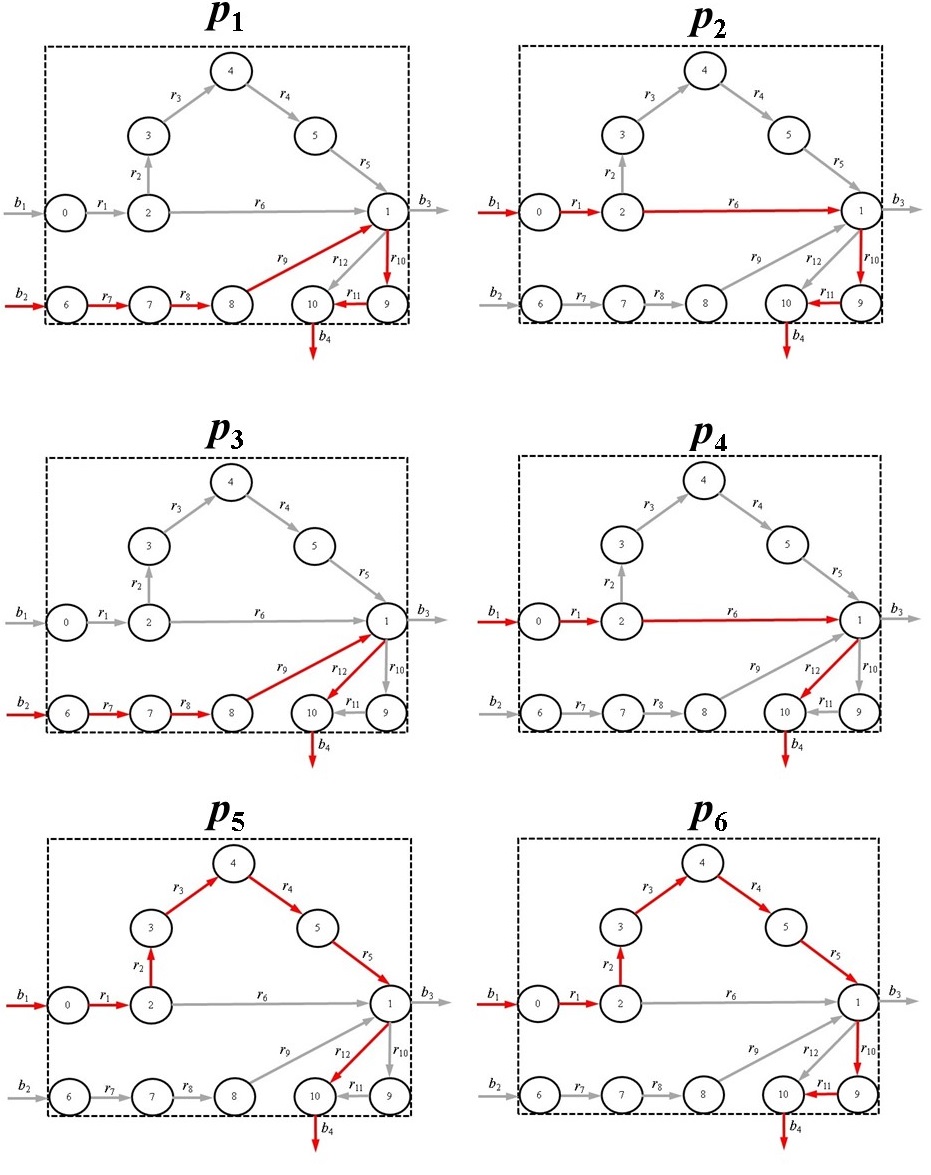

**Supplementary Fig. S2.** Elementary Modes obtained by metabolic pathway analysis in Klebsiella spp. bacteria. These EM show all the possible pathways of the system which main product is L-Glutamate.

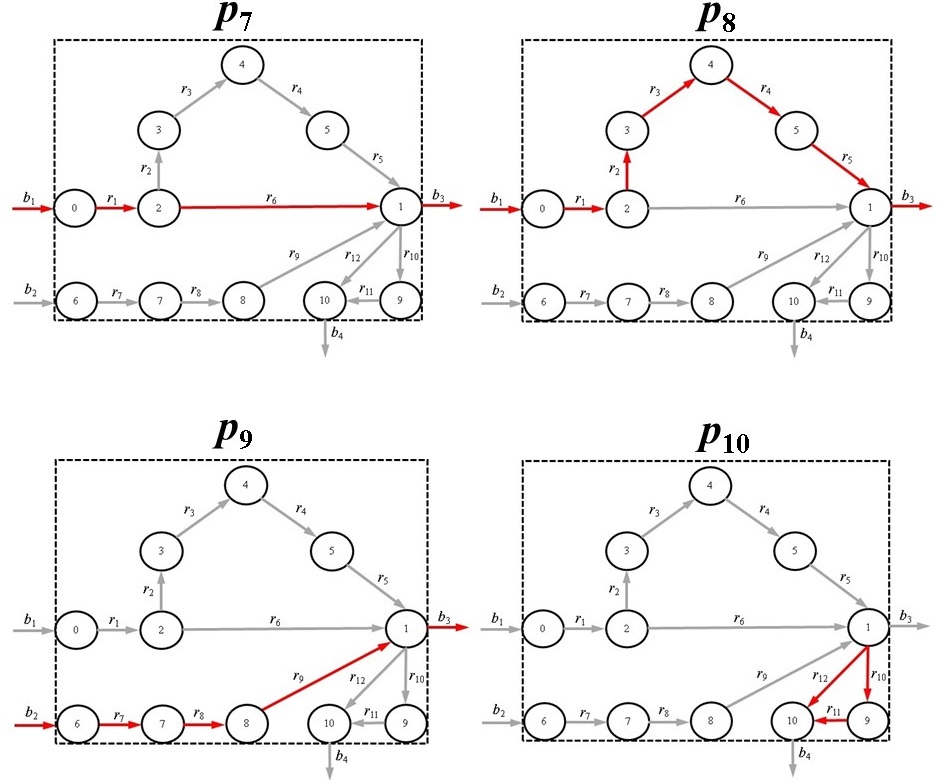

**Supplementary Fig. S3.** Elementary Modes obtained by metabolic pathway analysis in Klebsiella spp. bacteria. These EM show all the possible pathways of the system which main product is ammonia.
